# The bacterial SOS response promotes the expression of the transposase encoded by IS*CR* mobile genetic elements

**DOI:** 10.1101/2025.11.21.689736

**Authors:** Claire Lallement, Thomas Jové, Cécile Pasternak, Sandra Da Re, Marie-Cécile Ploy

## Abstract

Insertion sequences (IS) are widely involved in bacterial genomic plasticity by disrupting, adding, moving genomic sequences, or by activating or extinguishing gene expression. A specific family of IS, IS*CR* (for insertion sequence of Common Region), is thought to be involved in the dissemination of antibiotic resistance genes (ARG). While some IS*CR* members are commonly found in bacteria isolated in clinical settings and can contribute to downstream ARG expression, the mechanisms regulating the IS*CR*-encoded transposases expression have remained uncharacterized. Here, we investigated the expression of the transposase gene of ISCR1, ISCR2 and ISCR8, and its regulation in *Escherichia coli*. Using *in silico* analyses and *in vitro* experiments, we showed that expression levels were extremely low, as observed for most IS transposases. We further demonstrated the direct role of DNA damages and the key SOS response repressor, LexA, in controlling the activity of transposase promoter. These results provide evidence that the mobility of at least some IS*CR* elements may be promoted upon bacterial exposure to antibiotics inducing the SOS response.

**IMPORTANCE:** Mobile genetic elements are the most prevalent cause of antibiotic resistance emergence. Among these mobile elements, insertion sequences (IS) are well known to allow the dissemination of antibiotic resistance genes through the action of their tranposase. Here, we studied the regulation of the transposase expression in a specific family of IS, the IS*CR* family, some members of which are known to be involved in antibiotic resistance. Characterizing the regulation of transposase expression is an important starting point for understanding how these IS can contribute, through their movement, to the spread and expression of antibiotic resistance. This knowledge is necessary if we are to hope to prevent this spread one day.

## INTRODUCTION

Bacterial mobile genetic elements (MGEs) are key players in the dissemination of antibiotic resistance genes through horizontal gene transfer (1). Insertion sequences (IS) are the most widespread type of MGEs and encode all necessary material for their transposition (2). IS can differently impact the physiology of their host cells by moving, disrupting, or activating genes (3). IS elements are generally tightly regulated through diverse mechanisms acting at different levels, including transcriptional, translational and/or post-translational levels (reviewed in (4)). For instance, the transposition of IS*10* is controlled by the SOS response in *Escherichia coli* (5), IS*2* by the global regulator CRP (cAMP receptor protein) (6), and IS*50* by Hfq (7).

IS of the Insertion Sequence Common Region (IS*CR*) group are of clinical concern since they are commonly adjacent to multiple antibiotic resistance genes (ARGs) or pathogenicity regions (8). IS*CR* elements are related to the IS*91* family members with which they share several features. These include (i) a gene encoding a transposase of the HUH superfamily of single-strand nuclease (hereafter referred to as *rcr*, according to (9)) and (ii) two distinct ends that include sequences forming secondary structures, namely *ori*IS and *ter*IS, where the transposition process initiates and terminates, respectively (8, 10) (Fig. 1A). These specificities suggest that IS*CR* elements may transpose by a rolling-circle mechanism as presumed for IS*91*, and can therefore transpose adjacent genes (8, 11).

**Figure 1.**
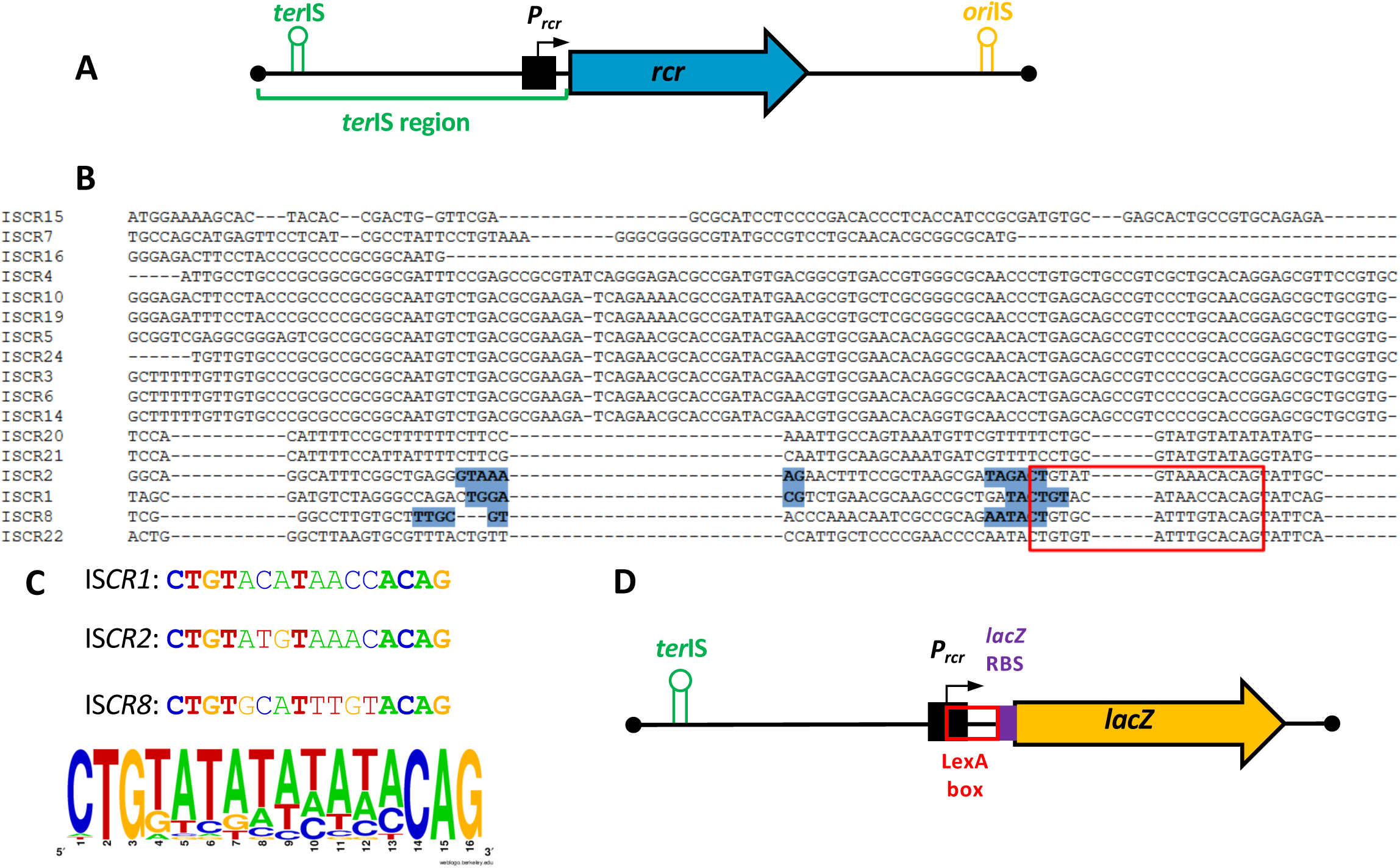
Structure of IS*CR* and alignment of their *ter*IS regions. **A. General structure of ISCR**. ISCR consists of a *rcr* transposase gene and its promoter P*_rcr_*, bound by two pairs of inverted repeats, namely *ter*IS and *oriI*S. **B. Alignment of *ter*IS regions.** Putative -35 and -10 σ^70^ elements of *rcr* promoters of interest are shown in bold and highlighted in blue. Putative LexA box are boxed in red. Aligned sequences include the *ter*IS region (up to to the *rcr* start codon) limited to 150 bp. Alignment made with Clustal Omega (https://www.ebi.ac.uk/jdispatcher/msa/clustalo) with default parameters. **C. Comparison of LexA operators in IS*CR1,* IS*CR2* and IS*CR8.*** Weblogo was used to generate the LexA box consensus sequence (Crooks *et al*, 2004) from a set of 28 experimentally verified *Escherichia coli* LexA binding sites (listed in Erill *et al*, 2003). Bases identical in each of the three predicted IS*CR* LexA binding sites and matching the LexA box logo are in bold. Each color represents a nucleotide (blue : cystosine, red : thymine, yellow : guanine, green : adenine) **D. Transcriptional fusion used to assess P*_rcr_* activity in this study.** Reporter gene *lacZ*, encoding for β-galactosidase, was fused in a translational fusion with native (P*_rcr_*), LexA-box mutated (P*_rcr_*L) or - 10 region mutated (P*_rcr_*M) promoters of IS*CR1*, IS*CR2* and IS*CR8*.

The prevailing IS*CR* element, namely IS*CR1*, is exclusively found downstream of the 3’ conserved region associated with class 1 integrons and displays a large diversity of adjacent genes, especially ARGs (8, 9). Moreover, IS*CR1* harbors two outwardly oriented promoters (P_OUT_) in its *ori*IS end (12–14), contributing to the expression of downstream ARGs (9). Additionally, two other IS*CR*s commonly found in the literature and databases are (i) IS*CR2*, which is mainly described adjacent to ARG within ICEs (Integrative Conjugative Element), especially the SXT element (15) and (ii) IS*CR8,* which is found as multiple copies in environmental bacterial plasmids (16).

Despite their potential importance in clinical settings, nothing is known about the regulation of the expression of IS*CR* elements.

In this study, we characterized, both *in silico* and *in vitro*, the key DNA motifs involved in the regulated expression of the *rcr* transposase-encoding genes of three members of the IS*CR* group (IS*CR1*, -*2* and -*8*), in *Escherichia coli*. Specifically, we demonstrate that the SOS response acts as a crucial pathway to tightly control the expression of these IS*CR* transposases, *via* the LexA repressor.

## RESULTS AND DISCUSSION

### LexA binds to the putative promoter region of three IS*CR* transposase (*rcr*) genes

To investigate the expression of IS*CR* transposase genes, we first analyzed *in silico* the *ter*IS region sequences encompassing the promoter region of the *rcr* gene (Figure 1A, B) of 17 non-redundant IS*CR* elements in *E. coli* (see Fig. S1). Interestingly, IS*CR1*, IS*CR2* and IS*CR8* comprised a sequence matching the CTGT-N_8_-ACAG β-/γ-proteobacteria LexA binding site consensus (LexA box) (17, 18), which overlaps with the -10 element of the predicted *rcr* transposase gene promoters (P*rcr*) (Fig. 1B, C), suggesting a potential regulation of these promoters by the repressor LexA. The LexA transcriptional factor is the master regulator of the SOS response, a global bacterial response to DNA damage (reviewed in (18)), that acts as a repressor of SOS regulon genes under normal conditions. Upon DNA damage and the formation of single-strand DNA (ssDNA), the SOS response is triggered, through the formation of a nucleoprotein filament ssDNA-RecA that stimulates LexA autoproteolysis and therefore the expression of SOS-regulated genes.

To confirm the functionality of the predicted LexA box in IS*CR1*, *-2* and *-8*, we performed electrophoretic mobility shift assay (EMSA) of their *rcr* promoter regions with purified *E. coli* LexA protein. We observed a DNA shift in the presence of the LexA protein for the three promoters P*_rcr_* (Fig. 2), showing that LexA specifically binds *in vitro* to the putative IS*CR* LexA box located in the native IS*CR1*, IS*CR2* and IS*CR8* P*_rcr_* promoter regions. Conversely, no shift was detected when the putative IS*CR* LexA boxes were mutated in one of the conserved terminal triplet (CTGT-N_8_-ACAG into CTGT-N_8_-A**<u>ACT;</u>** P*_rcr_*L) (Fig. 2). These results strongly suggest that LexA plays a role in regulating expression of the corresponding *rcr* transposase gene, and show that the sequences identified *in silico* are functional LexA boxes.

**Figure 2.**
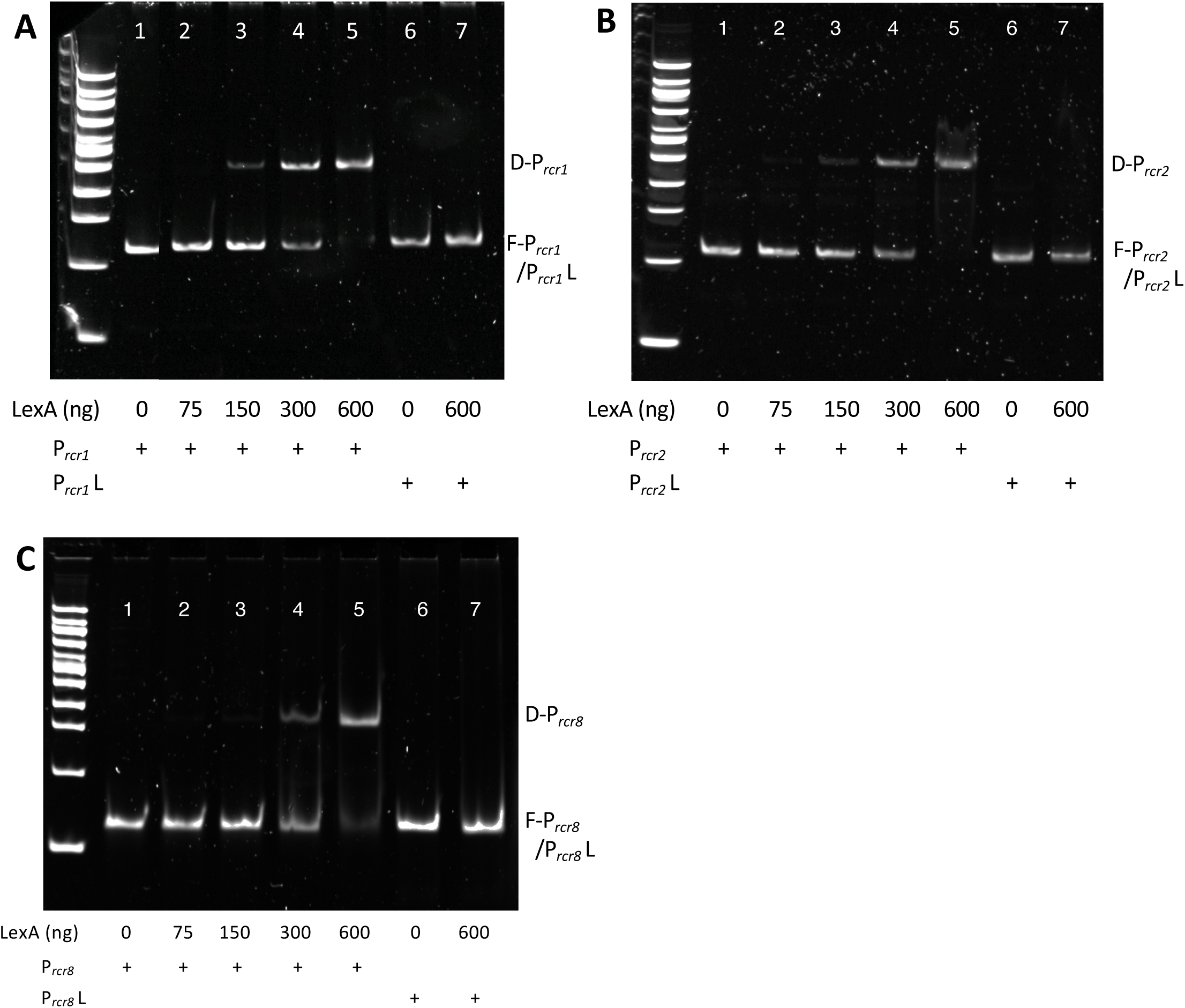
LexA binds to the *rcr* promoter region of IS*CR*1, IS*CR2* and IS*CR8*. Electrophoretic mobility shift assays performed with native *rcr* promoters (P*_rcr_* ; lanes 1-5) or LexA box-mutated *rcr* promoters (P*_rcr_* L; lanes 6-7) of *rcr1* (A), *rcr2* (B) or *rcr8* (C), in the presence or absence of purified LexA protein (amounts in nanograms are indicated). F, free DNA; D, delayed complex.

### The SOS response induces the IS*CR1,* IS*CR2* and IS*CR8* transposase expression

To characterize the SOS regulation of the *rcr* transposase genes *in vivo*, we constructed *lacZ* transcriptional fusions between the *lacZ* gene with its RBS and the *ter*IS regions of IS*CR1*, IS*CR2* and IS*CR8*, including the putative *rcr* promoter (P*_rcr_*) sequence and LexA-binding site (Fig. 1D). We also constructed these fusions containing a mutation of the putative -10 box of the P*_rcr_* promoters (P*_rcr_* M) or the mutation in the LexA box (P*_rcr_*L) (Table 1).

**Table 1:**
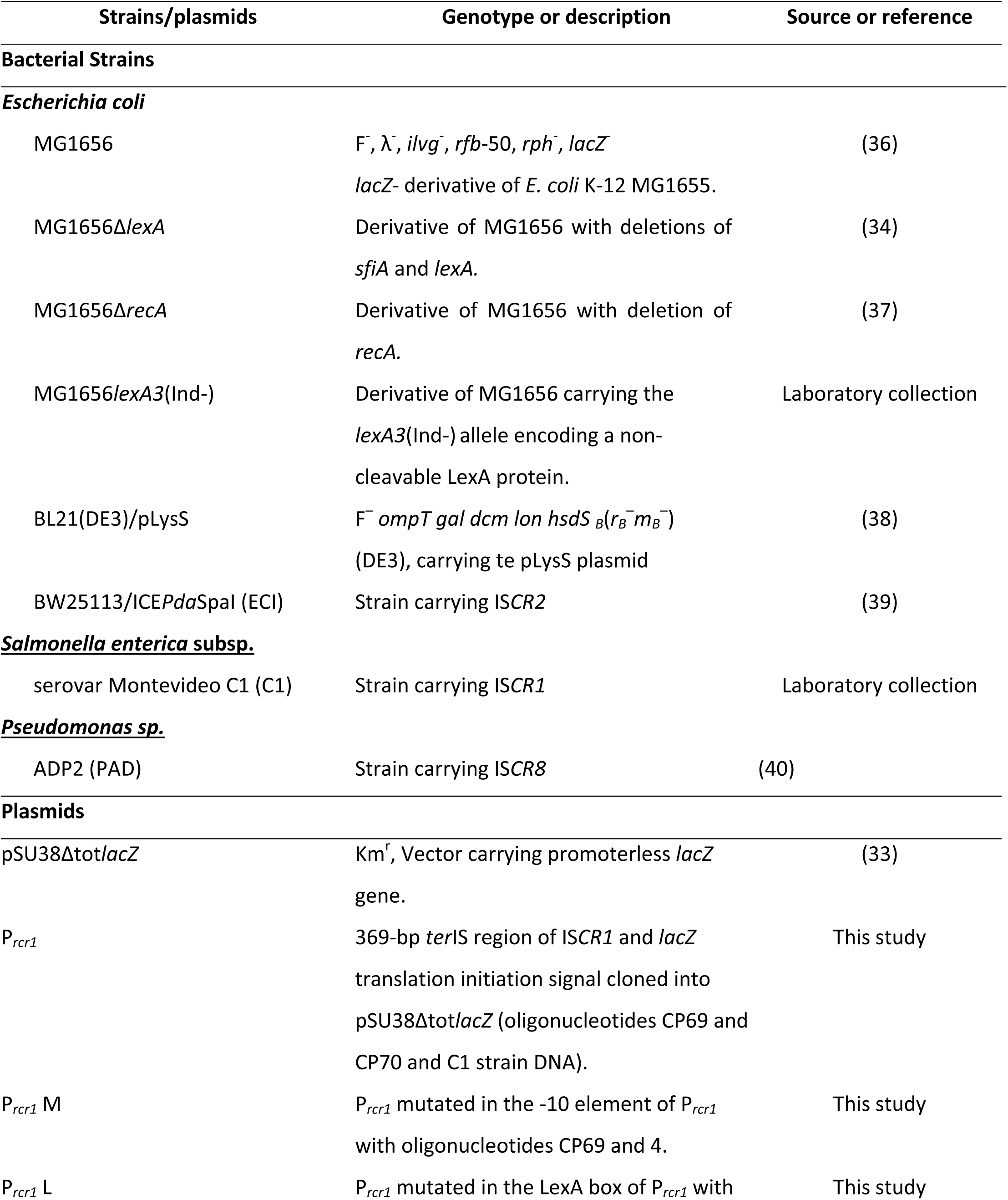
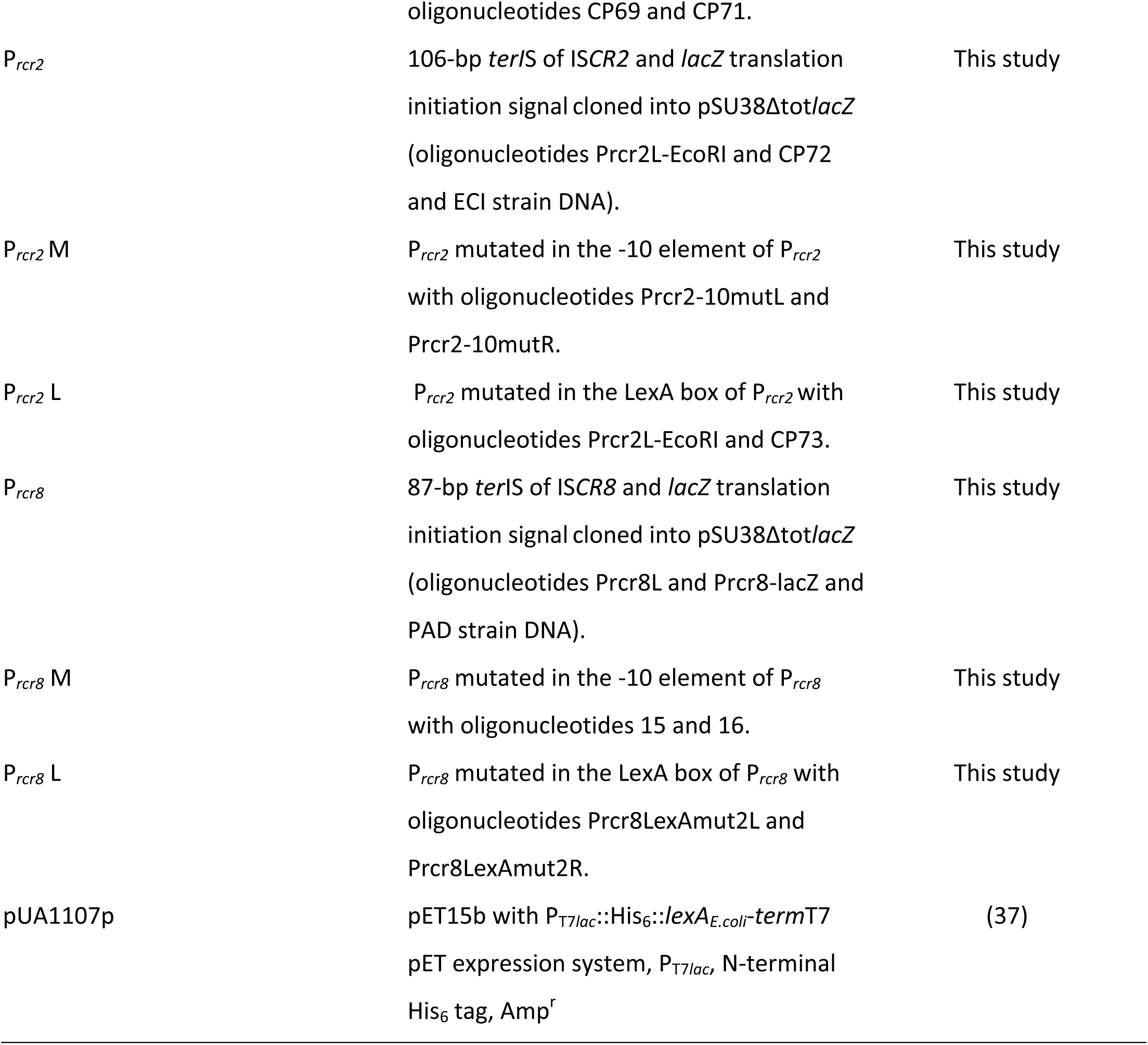
Bacterial strains and plasmids used in this study.

The resulting recombinant plasmids were introduced into the *E. coli* K-12 MG1656 strain, its Δ*lexA* (constitutive expression of LexA regulated promoters) and Δ*recA* (constitutive repression of LexA regulated promoters) derivatives, as well as in a *lexA3*(Ind-) mutant encoding for an uncleavable LexA protein (19). Measuring β-galactosidase activities showed very low activity for all 3 promoters in the WT strain (<5 uMiller), suggesting that Rcr transposases are not constitutively expressed (Fig. 3A-C). These results are consistent with the literature. Indeed, a weak promoter activity is a feature shared by many promoters from transposase genes, including those from IS*21* (20), IS*30* (21), IS*911* (22) and IS*200* (23), to limit their impact on bacterial fitness. We observed significant increase in β-galactosidase activity in the *lexA*-deleted strain, with varying strength for the three native promoters P*_rcr1_,* P*_rcr2_* and P*_rcr8_*(18.2-, 1.6- and 4.1-fold respectively; Fig. 3A-C and Table S1). However, we did not observe a strong effect in conditions where the SOS response could not be induced (strains Δ*recA* and *lexA3*(Ind-))-, basal P*_rcr_* activities being very low (<5 uMiller) in the WT strain (Fig. 3D). Overall, the results are consistent with the potential regulation of P*_rcr1_*, P*_rcr2_*, and P*_rcr8_* by LexA. To further validate these results, we measured the β-galactosidases activities from P*_rcr_*L mutated promoters (no binding of LexA). As expected, the mutation of the LexA-binding site led to a 48.5-, 14.5- and 10.6- fold increase in β-galactosidase activities for P*_rcr1_*, P*_rcr2_* and P*_rcr8_*, respectively (Fig. 3A-C and Table S1). These results demonstrate the involvement of LexA protein in the regulation of IS*CR1*, *-2* and *-8 rcr* promoters.

**Figure 3:**
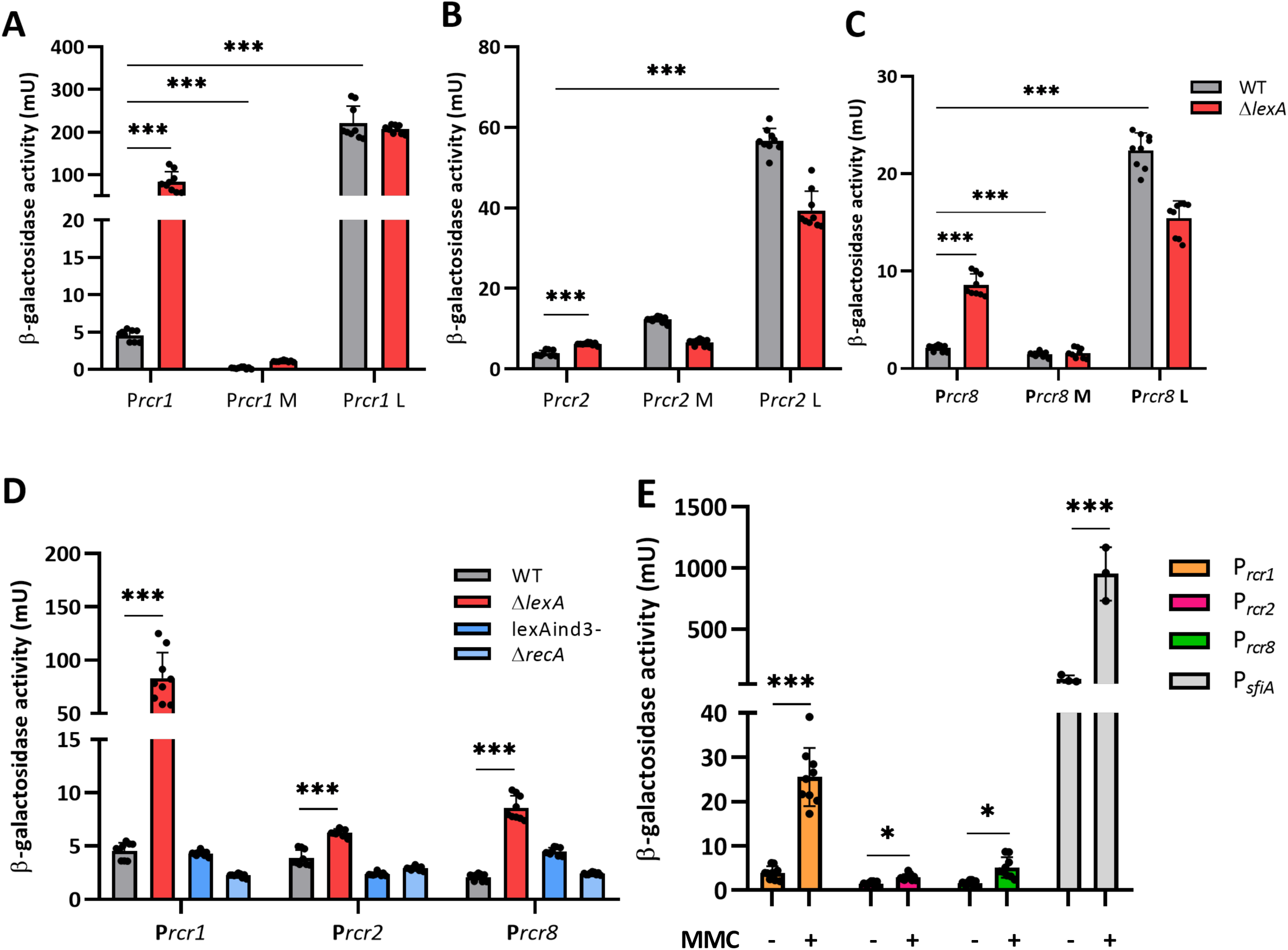
The SOS response controls the expression of IS*CR1*, IS*CR2* and I*SCR8* transposase *rcr* genes A-D. LexA represses expression of the transposase genes. β-galactosidase activities (in Miller units mU) were measured in the *E. coli* MG1656 reference strain (grey) and its Δ*lexA* (red), Δ*recA* (light blue) and *lexAind3-* (non-autoproteolytic LexA; turquoise)) derivatives. These strains carry a plasmid containing a *lacZ* transcriptional fusion with the *rcr* promoter of IS*CR1* **(A, D)**, IS*CR2* **(B, D)** or IS*CR8* **(C, D)** in either the native (P*_rcr_*), LexA-box mutated (P*_rcr_*L) or -10 region mutated (P*_rcr_*M) promoter configuration as indicated. Experiments were performed 3 times in technical triplicates. Statistical significance was determined by unpaired two-tailed t tests (*** *p*-value <0.01). **E. The IS*CR* transposase genes expression is induced by SOS response inducers.** β-galactosidase activities were measured in the *E. coli* MG1656 reference strain carrying a plasmid containing a *lacZ* transcriptional fusion with P*_rcr1_*(orange), P*_rcr2_* (red), P*_rcr3_* (green) or PsfiA (light grey; *E. coli SOS* induced promoter), after treatment (+) or not (-) with mitomycin C (MMC), an SOS inducing agent. All experiments were performed 3 times in technical triplicates. Statistical significance was determined by unpaired two-tailed t tests (*** *p*-value <0.01, * *p*-value <0.05).

To confirm the SOS induction, we measured P*_rcr1_*, P*_rcr2_* and P*_rcr8_* activity after treatment with mitomycin C (MMC), a DNA-damaging molecule known to induce the SOS response in *E. coli* (24, 25). As shown in Fig.3E, treatment with MMC induced a significant and strong increase (7-fold) in β-galactosidase activity for P*_rcr1_* and milder ones for P*_rcr2_* and P*_rcr8_* (2- and 3.2-fold respectively; Table S1).

The intrinsic activity of the three P*_rcr_* promoters was then analyzed to validate the putative promoter sequences identified *in silico* (Fig. 1B) by introducing mutations into the -10 region of the P*_rcr_* promoters (P*_rcr_* M ; see Material and Methods). As anticipated from the very weak activity of the native promoters, P*_rcr_* M mutation almost completely abolished activity of P*_rcr1_* and P*_rcr8_* in the Δ*lexA* background only, demonstrating the existence of an active promoter tightly repressed by LexA (Fig. 3A, 3C and Table S1). Surprisingly, for P*_rcr2_* mutated in the -10 region, we obtained a 3.1- fold higher β-galactosidase activity compared to P_rcr2_, in the wild type strain. We did not observe such an increase in the Δ*lexA* strain, (Fig. 3B). These results suggest either that the P*_rcr2_* promoter has not been correctly defined by the *in silico* approach, or that the mutations performed to inactivate this promoter indeed disrupted its activity but allowed, at the same time, the recognition of a slightly more active promoter. We performed another *in silico* analysis confirming that the introduction of the mutations in the initial putative -10 region of P*_rcr2_* allowed the identification of a new P*_rcr2_* promoter upstream of the initial one (Fig. S2). This could explain the increased activity for P*_rcr2_*M in wild-type background.

### Implications for IS*CR* mobility

Although it is well known that the expression of genes involved in the mobility in many diverse MGEs, like prophages, plasmids, pathogenic islands, gene cassettes from integrons (reviewed in (26)) and ICE (24), is controlled by the SOS response through LexA repression, only a few articles in the literature report the contribution of the host SOS system in the regulation of IS transposable elements. The SOS response has been involved in the UV-induced transposition of IS*10* (5) and possibly in the transposition of Tn*5* (IS*50*) in *E. coli* (27), but controversy still exists regarding the role of RecA and LexA in regulating Tn*5* (28, 29). Here, we demonstrate that the expression of the transposases of ISCR1 and ISCR8 is directly and tightly regulated by the repressor of the SOS response, LexA. Although LexA can specifically bind to the *rcr* promoter region of IS*CR2 in vitro*, P*_rcr2_* activity remains extremely low (below 5 miller units) even when induced with SOS-inducing agents or in the Δ*lexA* mutant (less than 2-fold induction, Fig. 3 B and E). The *rcr2* gene being frequently associated with the SXT ICE (8), we can hypothesise that IS*CR2* would no longer be mobile and would depend solely on SXT ICE mobility to be transmitted.

Intriguingly, IS*CR1* appears to be restricted to β-/γ-proteobacteria (Lallement et al., 2018) *i.e.* hosts displaying a canonical SOS response. This highlights an interesting strategy for the IS*CR1* element: using a host-dependent negative regulation not only allows a low fitness cost, but also to carry solely the gene required for mobility that will only be expressed under drastic conditions via host regulation. Furthermore, the transposases encoded by IS*CR* elements are members of the HUH family of endonucleases that use as a substrate a ssDNA known to trigger the SOS response. Thereby, the availability of the ssDNA substrates for IS*CR* transposase may be sensed as a signal to stimulate the transposase expression through the induction of the SOS response, ultimately promoting their transposition. Although the transposition mechanism of the IS*CR* elements has not been experimentally proven, the present study demonstrates that under DNA-damaging conditions, the transposase genes of some IS*CR* elements can be expressed, which could contribute to the spread of IS*CR* and adjacent AR genes. However, as their level of expression remains extremely low, even when de-repressed, IS*CR* 1, 2 and 8 elements, may mostly rely on the MGE carrying them, such as plasmids (for IS*CR1* and ISCR8) or ICE (for IS*CR2*) to spread.

This study demonstrates a direct regulation of the expression of the transposase gene by the SOS response and its repressor LexA, in several members of the ISCR/IS91 family of bacterial IS elements. As we witness an ever-increasing spread of antibiotic resistance globally, understanding the mechanisms that govern this spread is an obvious necessity. In this context, our findings reveal the importance of the host SOS response in the mobility of IS*CR*, and offer evidence that, by relying on host regulatory mechanisms, some IS elements exhibit β-/γ-proteobacteria host specificity, and can carry minimal genetic material for their own regulation.

## MATERIAL AND METHODS

### Bioinformatic analysis of IS*CR* elements

Seventeen IS*CR* members considered in this study were recovered from the initial list published by Toleman *et al.* (2006), completed by the current updated list from the Galileo database (https://galileoamr.arcbio.com/mara/), and curated to discard IS*CR* elements sharing more than 98% nucleotide identity (8, 30). When available, we aligned the sequences upstream of the transposase START codon up to 150 bp long. Putative σ^70^ promoters were predicted by BPROM (www.softberry.com (31)). Alignments were performed using Clustal Omega (https://www.ebi.ac.uk/jdispatcher/msa/clustalo) (32) with default parameters. Sequences and accession number of all IS*CR* used for this alignment are in Supplementary data files.

### Bacterial Strains and Plasmids

Bacterial strains and plasmids are listed in Table 1. Bacterial strains were grown in Lysogeny Broth (LB) at 37°C. Medium was supplemented with kanamycin (25 µg/mL) when required. The expression of *E. coli* His_6_::LexA protein in *E. coli* BL21(DE3)/pLysS was induced by adding 1 mM isopropyl- β-D-thiogalactopyranoside (IPTG) to the medium.

### DNA manipulations

Plasmid DNA was extracted from *E. coli* using the Wizard^®^ Plus SV Minipreps DNA Purification Kit (Promega, Madison, WI, United States), and genomic DNA using the SaMag Bacterial DNA Extraction Kit (Sacace, Biotechnologies). PCR reactions were carried out with Phusion^®^ DNA Polymerase (Thermo Fisher Scientific) to amplify fragments subsequently used for cloning, or with the Taq’Ozyme Purple Mix 2 (Ozyme) to screen transformants. Oligonucleotides used in this study are listed in Table S2. PCR products were loaded and visualized on 1.6% agarose gel, extracted and purified with the Wizard^®^ ^SV^ Gel (Promega, Madison, WI, United States) or PCR Clean-Up kit (Macherey-Nagel).

### Plasmid Constructions

Transcriptional *lacZ* fusions were constructed in the pSU38Δtot*lacZ* reporter plasmid (33). The *rcr* promoter region amplified from bacterial strains known to carry the corresponding IS*CR* was cloned into the EcoRI/BamHI unique restriction sites of pSU38Δtot*lacZ* (see Table 1). Mutations in LexA boxes were performed by PCR assembly to convert the LexA box essential triplet “CAG” into “ACT”, as previously done (34). The P*_rcr_* promoters were mutated by PCR assembly in their putative -10 elements: from TACTGT to **<u>CG</u>**CTGT (IS*CR1*; P*_rcr1_* M), TAGACT to **<u>CG</u>**GACT (IS*CR2*; P*_rcr2_* M) and TACTGT to **<u>GC</u>**CTGT (IS*CR8*; P*_rcr8_* M).

### β-galactosidases assays

β-galactosidase assays were performed in the *E. coli* MG1656 strain from at least three independent assays and three technical replicates for each construct as previously described at 37°C (33). The SOS response was induced with the addition of mitomycin C (MMC) 1.6 µg/mL in the culture one hour before reaching 0.6-0.8 OD_600_.

### Electrophoretic Mobility Shift Assay (EMSA)

Over-expression and purification of the *E. coli* His_6_::LexA protein was performed as described previously (35). EMSA probes, with natural (P*_rcr_*) or mutated LexA binding site (P*_rcr_* L), were amplified by PCR from the C1 (IS*CR1*) or ECI (IS*CR2*) bacterial strains (Table 1) using oligonucleotide pairs ISCR1-GS-L/ISCR1-GS-R, ISCR1-lexAmutL/ISCR1-lexAmutL, Prcr2L-EcoRI/ISCR2_GS_R and Prcr2-LexAmutL/Prcr2-LexAmutR for probes P*_rcr1,_*P*_rcr1_* L (218bp), P*_rcr2_* and P*_rcr1_*L (209bp) respectively (Table S2). EMSA probes were purified on agarose gel 1.6%. For P*_rcr8_*, the WT and mutated probes were ordered from Merck as bottom and top strands (C129+CP130 and CP131+CP132 respectively; Table S2) that were assembled prior EMSA experiment.

Increasing quantities of recombinant His_6_::LexA protein (0 to 600 ng) were incubated on ice for 20 min with 40 ng of DNA probes in a final volume of 10 µL of binding buffer (10 mM HEPES pH 8, 10 mM Tris-HCl pH 8, 50 mM KCl, 1 mM EDTA). Then, 2 µL of 6X EMSA gel-loading solution (EMSA Kit, Molecular Probes BioRad) were added to the reaction mixture. Samples were separated in 6% non-denaturing Tris-glycine polyacrylamide gels and visualised with SYBR® Green following the manufacturer’s protocol (EMSA Kit, Molecular Probes BioRad).

### Statistical analysis

Statistical analysis was performed using the t-test with two unpaired groups.

## ACKNOWLEDGMENTS

This paper is dedicated to the Cécile Pasternak, an inspiring scientist, motivating mentor, and dear colleague. Cécile touched the lives of countless numbers of students and colleagues. She was an outstanding microbiologist and a real well of science who taught us so much. We miss our conversations, scientific and otherwise. Cécile passed away on 23 January 2023, and our lives now lack such a kind and inspiring person. Thank you Cécile.

This work was supported by grants from Ministère de l’Enseignement Supérieur et de la Recherche and Institut National de la Santé et de la Recherche Médicale (Inserm). CL gratefully acknowledges the Ministère de l’Enseignement Supérieur, de la Recherche et de l’Innovation (MESRI), and the Fonds Européen de Développement Régional (FEDER) for her doctoral training grant. The funders had no role in study design, data collection and interpretation, or the decision to submit the work for publication.

The authors would like to acknowledge Yohann Lacotte for helpful discussions as well as Vincent Burrus, Bénédicte Michel, Marion Devers, Fabrice Martin and Bao Ton-Hoang for their gift of strains, and Anna Thieulot for experiments done during her master 1 internship. We thank Emmanuelle Botté from ManuScribes for professional editing of the manuscript.

## SUPPLEMENTAL MATERIAL

**Fig. S1:** Sequences and accession number of IS*CR* family members used for Figure 1B alignment.

**Fig. S2:** Mutation in -10 region generates an alternative promoter in P*_rcr2_*.

**Table S1:** β-galactosidase activities from P*_rcr1_*, P*_rcr2_* and P*_rcr8_* promoters and their derivatives. Table S2: Oligonucleotides used in this study.

